# Intestinal Gluconeogenesis Regulates Brown and White Adipose Tissues Functions in mice

**DOI:** 10.1101/2021.10.25.465675

**Authors:** Justine Vily-Petit, Maud Soty-Roca, Marine Silva, Manon Micoud, Clara Bron, Margaux Raffin, Daniel Beiroa, Rubén Nogueiras, Damien Roussel, Amandine Gautier-Stein, Fabienne Rajas, Daniela Cota, Gilles Mithieux

## Abstract

**Objective:** Intestinal gluconeogenesis, via the initiation of a gut-brain nervous circuit, accounts for the metabolic benefits linked to dietary proteins or fermentable fibre in rodents and has been positively correlated with the rapid amelioration of body weight after gastric bypass surgery in obese humans. In particular, the activation of intestinal gluconeogenesis moderates the development of hepatic steatosis accompanying obesity. In this study, we investigated the specific effects of intestinal gluconeogenesis on adipose tissue metabolism, independently of its induction by nutritional manipulation.

**Methods:** We used two transgenic mouse models of suppression or overexpression of G6PC, the catalytic subunit of glucose-6 phosphatase, the key enzyme of endogenous glucose production, specifically in the intestine.

**Results:** Under a hypercaloric diet, mice with a genetic overexpression of intestinal gluconeogenesis showed a lower adiposity and higher thermogenic capacities than wild-type mice, featuring marked browning of white adipose tissue and prevention of the whitening of brown adipose tissue. Suppression of sympathetic nervous signalling in brown adipose tissue impairs the activation of thermogenesis. Conversely, mice with genetic suppression of intestinal gluconeogenesis exhibit an increase in adiposity under standard diet, associated with a decreased expression of markers of thermogenesis in both the brown and white adipose tissues.

**Conclusion:** **I**ntestinal gluconeogenesis is sufficient in itself to activate the sympathetic nervous system and prevent the expansion and the metabolic alterations of brown and white adipose tissues metabolism under high calorie diet, thus preventing the development of obesity. These data increase knowledge of the mechanisms of weight reduction in gastric bypass surgery and pave the way of new approaches to prevent or cure obesity.

## 1. INTRODUCTION

The world is faced with an exponential increase in the prevalence of obesity. Unfortunately, the development of type 2 diabetes, which often accompanies the development of obesity, poses serious health problems, as retinopathy, nephropathy, neuropathy, cardiovascular diseases and various cancer types [1].

Adipose tissue, through its functions of energy storage and secretion of numerous hormones (the adipokines), is a crucial organ in energy balance. In mammals, the white adipose tissue (WAT) is mainly dedicated to lipid storage while the brown adipose tissue (BAT) is specialized in heat production. Both BAT and WAT can respond quickly to nutrient excess through adipocyte hypertrophy and hyperplasia. Adipose tissue remodeling is accelerated during the development of obesity, which involves metabolic alterations such as deregulation of fatty acid fluxes and dyslipidemia, mitochondrial dysfunctions and changes in adipokine synthesis. This leads to insulin resistance, chronic inflammation, fibrosis and even cell death [2]. In the early stages of obesity development, the WAT is a preferential storage site for excessive circulating lipids and is the first to undergo deleterious changes. However, the BAT is also recruited for lipid storage, and gradually loses its thermogenic capacity. This defect in turn contributes to the appearance of type-2 diabetes and its associated complications [3]. What is more, compelling data strongly suggest that deregulated adipose tissue metabolism, via inflammation, could be an initiating factor in the development of type-2 diabetes [4].

In this context, it is noteworthy that intestinal gluconeogenesis (IGN) positively interferes with the control of glucose and energy homeostasis to exert anti-diabetic and anti-obesity effects. More specifically, glucose released in the portal vein is sensed by a glucose receptor present in the gastrointestinal neural system (sodium-glucose co-transporter-3, SGLT3), which initiates a nervous signal to the hypothalamic nuclei regulating energy homeostasis [5–7]. The metabolic benefits previously observed in response to various nutritional manipulations in mice such as upon protein- or fibre-enriched diets, were strongly suggested to depend on the induction of IGN [8–11]. Moreover, IGN has been positively correlated with the rapid ameliorations in energy homeostasis after gastric bypass surgery, both in mice and in obese humans [12–14]. Thus, IGN participates to the prevention of body weight gain, the reduction in fat mass expansion and the improvement in glycaemic control under high calorie diet [7,8,14,15]. Using a mouse model of constitutive overexpression of glucose-6-phosphatase (G6Pase) in the intestine, we recently reported that the induction of IGN *per se* prevents hepatic steatosis under high calorie diet [16]. However, it remained to know whether there is a beneficial effect of IGN on adipose tissue physiology and the underlying mechanisms.

In this work, we deciphered the mechanisms evoked by IGN in the regulation of both WAT and BAT function. We studied: **1/** whether the genetic induction of IGN could protect adipose tissue metabolism in a deleterious nutritional context (high fat-high sucrose (HF-HS) diet); **2/** whether the lack of IGN could be sufficient to lead to alterations in adipose metabolism in the context of a standard (starch-based) diet. We report that IGN, via the recruitment of the local sympathetic system, regulates key functions of adipose tissue metabolism.

## 2. METHODS

### 2.1. Animals, diets and ethical statement

We used two transgenic mouse models of suppression (I.G6pc^-/-^ mice [11]) or overexpression (I.G6pc^overexp^ mice [16]) of IGN by targeting *G6pc1*, the catalytic subunit of G6Pase that is the last enzyme operating before glucose release. Wild-type C57Bl6/J (WT, Charles River Laboratories, France) mice were used as control mice. All the mice were housed in the animal facility of Lyon 1 University (“Animalerie Lyon Est Conventionnelle and Specific Pathogen Free”) under controlled temperature (22°C) conditions, with a 12-h light/dark cycle, and *ad libitum* access to water and food. I.G6pc^-/-^ mice received standard chow diet and I.G6pc^overexp^ mice received HF-HS (5-months) or standard chow diet [16]. This study was conducted in males aged of 13 weeks (standard diet) or 8-months (HF-HS diet).

All procedures were performed in accordance with the principles and guidelines established by the European Convention for the protection of Laboratory Animals. The regional animal care committee (CEEA-55, University Lyon I, France) approved all the experiments herein.

### 2.2. Histological and immunostaining analysis

For histological analyses of adipose tissues, formalin-fixed and paraffin-embedded tissues were cut in 4 μm thick sections and stained with hematoxylin and eosin staining, Masson’s Trichrome staining, or immunostaining. For immunostaining, slices were incubated overnight at 4°C with primary antibodies produced in rabbits directed towards PRDM16 (PR domain containing 16) and UCP1 (Uncoupling Protein 1) (see supplementary material). The slices were then incubated for 30 minutes at room temperature with a secondary antibody coupled to Horseradish peroxidase and revealed using a 3,3’-diaminobenzidine solution. The nuclei were labelled with haematoxylin. The slices were observed using an inverted microscope (Nikon Eclipse TS2R).

For the slices of epididymal (eWAT) and subcutaneous (scWAT) white adipose tissue the number of adipocytes per mm² were determined using ImageJ software (n=6; 4-15 pictures per mice). For the slices of BAT, the area occupied by the lipid droplets was determined using ImageJ software (n=6; 4-15 pictures per mice).

### 2.3. Electron Microscopy

A piece (1mm^3^) of eWAT and BAT was finely cut and added to a solution of glutaraldehyde 4% and cacodylate buffer 0.2M (CIQLE platform, Université Lyon 1, SFR Santé Lyon-Est). Sections were observed with a transmission electron microscope JEOL-1400JEM (Tokyo, Japan) operating at 80kV equipped with a camera Orius-1000 gatan and Digital Micrograph. The number and the surface area of mitochondria were determined relatively to the cytoplasm area (excluding the lipid droplet) using ImageJ software (n=4; 10 pictures per mice).

### 2.4. Magnetic resonance imaging analysis

The measurement of body composition was using nuclear magnetic resonance imaging (MRI, Whole Body Composition Analyzer; EchoMRI; Houston, TX), as previously described [17,18] (n=6-7).

### 2.5. Temperature measurements

Rectal and skin temperature surrounding BAT were measured and analyzed as previously described [17,18]. Briefly, body temperature was recorded with a rectal probe connected to a digital thermometer (BAT-12 Microprobe-Thermometer; Physitemp, Clifton, NJ). Skin temperature surrounding BAT was measured with an infrared camera (B335-Thermal Imaging Camera; FLIR; West Malling, Kent, UK) [20,21] and analyzed with FLIR-Tools-Software (n=6-7; 2 pictures per mice).

### 2.6. Chemical denervation of the sympathetic nervous system and cold exposure

Chemical denervation of BAT sympathetic nerve endings was performed in I.G6pc^overexp^ and WT mice fed standard diet [19] under isoflurane anaesthesia and ketoprofen (1mg/kg) analgesia. This technique using several injections of 6-hydroxydopamine (6-OHDA), a neurotoxin selective for sympathetic neurons, permits a specific denervation, while keeping the sensory fibres intact. The effects of sympathetic denervation were compared to those obtained after vehicle injection (0.15mol/L NaCl, 1% ascorbic acid).

Briefly, the two lobes of brown interscapular adipose tissue were exposed through a midline skin incision along the upper dorsal surface and gently separated from the skin with surgical forceps. Then, several injections of 6-OHDA (Sigma-Aldrich) were performed directly into each lobe of the interscapular BAT. For each lobe, 10µL (6-OHDA-10mg/mL) was injected in several times using a Hamilton syringe. The skin incision was then closed with several surgical stitches.

After 10 days of post-operative recovery, I.G6pc^overexp^ and WT mice were fasted for 5 hours, then re-fed and exposed to a cold temperature of 6°C for 4-hours. Rectal temperature and food intake were measured.

### 2.7. Measurement of mitochondria respiratory capacity

Fresh sample of BAT was homogenized with Dounce in a solution (250mM Sucrose, 1mM EGTA, 20mM Tris-base, 1% BSA, pH 7.3). Mitochondrion-enriched homogenate was obtained by successive centrifugations (2x(1000g; 10min), 2x(10 000g; 10 min)). The determinations were performed at 37°C on 1mg/mL of protein in 200µL of solution composed of 120 mM KCl, 5mM KH2PO4, 1mM EGTA, 2mM MgCl2, 3mM Hepes, 0.3% BSA, pH 7.4. Oxygen consumption was measured in a glass cell fitted with a Clark oxygen electrode (Rank Brothers Ltd) following the addition of different substrates, inhibitor, and uncoupler of the oxidative phosphorylation: Pyruvate/Malate (5mM/2,5mM), ADP (Adenosine-diphosphate, 1mM), Succinate (5mM), Oligomycin (2,5µg/mL), p-trifluoromethoxy-carbonyl-cyanide-phenyl hydrazone (FCCP; 5µM).

### 2.8. Gene expression analysis

Pieces of 150 mg of frozen eWAT or 20 mg of frozen BAT were homogenized using FastPrep® in Trizol reagent and total RNAs were isolated according to the manufacturer protocol (Invitrogen Life Technologies). Reverse transcription was done on 1µg of mRNA using the Qiagen QuantiTect Reverse Transcription kit. Real-time qPCRs were performed using sequence-specific primers supplied by Eurogentec (see supplementary material) with SsoAdvancedTM Universal SYBR® Green Supermix in a CFX-ConnectTM Real-Time System (Biorad).

The expression of mRNAs was normalized to the L32 (for eWAT) and cyclophilin (for scWAT and BAT) expression. The expression of target mRNAs was calculated using the 2-ΔΔCt method and expressed relatively to WT controls.

### 2.9. Western blot analysis

Cell extracts from eWAT were lysed in standard lysis buffer (20mM Tris-HCl, pH 8, 138mM NaCl, 1% NP40, 2.7mM KCl, 1 mM MgCl2, 5%glycerol, 5mM EDTA, 1mM Na3VO4, 20mM NaF, 1 mM DTT, 1% protease inhibitors), and homogenized using FastPrep®. Proteins were assayed in triplicate with Pierce™ BCA Protein Assay Kit (Thermo Fisher Scientific).

#### Denaturing conditions

Aliquots of 30 μg of proteins, denatured in buffer (20% glycerol, 10% β-mercaptoethanol, 10% SDS, 62.5mM Tris) were analysed from 4%, 9% or 12% -SDS polyacrylamide gel electrophoresis and transferred to PVDF Immobilon membranes (Biorad). After 1h-saturation in TBS/0.2% Tween/2% milk (RT), the membranes were probed (overnight at 4°C) with rabbit or mouse antibodies diluted in TBS/0.2% Tween/2% milk, against different proteins of interest, which names, references and dilutions are specified in the supplemental methods. Then, membranes were rinsed three-times in TBS/0.2% Tween for 10min and incubated for 1h with goat secondary anti-rabbit or anti-mouse IgG linked to peroxidase (dilution 1:5,000; Biorad) in TBS/0.2% Tween/2% milk. Membranes were rinsed again and exposed to ClarityTM Western ECL Substrate (Biorad). The intensity of the spots was determined by densitometry with ChemiDoc Software (Biorad) and analysed using the Image LabTM software (Biorad). Quantification of the αβ-tubulin, β-Actin or whole protein levels (using the stain free protocol provided by Biorad) was used for normalization. Stain-Free technology enabled fluorescent visualization of 1-D SDS PAGE gels and corresponding blots. The relative amount of total protein in each lane on the blot was calculated and used for quantitation normalization. Western blots were trimmed on the sides to show 3-4 representative samples of each experimental condition.

#### Non-denaturing conditions

To analyse the different conformations of adiponectin, aliquots of 30µg of protein were prepared in native Tris glycine sample buffer and analysed from NuPAGE 3-8% Tris-Acetate polyacrylamide gel electrophoresis in a tris-glycine native running buffer (Thermo Fisher Scientific). After 2-hours of migration at 130 V, proteins were transferred to PVDF Immobilon membranes (Biorad). After 1h-saturation in TBS/0.2% Tween/2% milk (RT), the membranes were probed (overnight at 4°C) with rabbit antibodies diluted in TBS/0.2% Tween/ 2% milk (see supplementary material). Then, membranes were rinsed three-times in TBS/0.2% Tween for 10min and incubated for 1h with goat secondary anti-rabbit IgG linked to peroxidase (dilution 1:5 000, Biorad) in TBS/0.2% Tween/2% milk. Membranes were rinsed again and exposed to ClarityTM Western ECL Substrate (Biorad). The intensity of the spots was determined by densitometry with ChemiDoc Software (Biorad) and analysed using the Image LabTM software (Biorad). The results were expressed as a ratio between the HMW and MMW forms of adiponectin (High molecular weight / Medium molecular weight).

### 2.10. Statistical analysis

All data are presented as mean ± SEM (standard error of the mean). Two-group and multi-group comparisons were analysed using unpaired *t test* or ANOVA-2 followed by Bonferroni post-hoc test, respectively. Values were considered significant at * p□<□0.05; ** p<0.01 and *** p<0.001. Statistical details and exact value of “n” can be found in the figure legends. Statistical analyses were performed with GraphPad Prism 9 software.

## 3. RESULTS

### 3.1. Intestinal gluconeogenesis prevents the development of fat mass in mice fed a high fat-high sucrose diet

In agreement with our previous studies reporting the protective effect of IGN overexpression against the development of obesity, MRI data indicated that I.G6pc^overexp^ mice had a lower fat mass than WT mice fed a HF-HS diet (**Fig. 1a**). Consistently with this observation, hematoxylin-stained eWAT and scWAT sections showed an increased number of adipocytes per mm² in I.G6pc^overexp^ mice compared to WT mice (**Fig. 1b**), highlighting adipocyte hypotrophy. Moreover, adipocytes in the BAT of I.G6pc^overexp^ contained smaller and greater number of lipid droplets than WT mice. This translates into a smaller lipid droplets area (**Fig. 1b**). In line with this reduction in lipid storage, the expression levels of *Perilipin*, which is the major lipid droplet coat protein in mature adipocytes was decreased in eWAT and BAT of I.G6pc^overexp^ mice as compared to WT mice (**Fig. 1c**).

**Figure 1:**
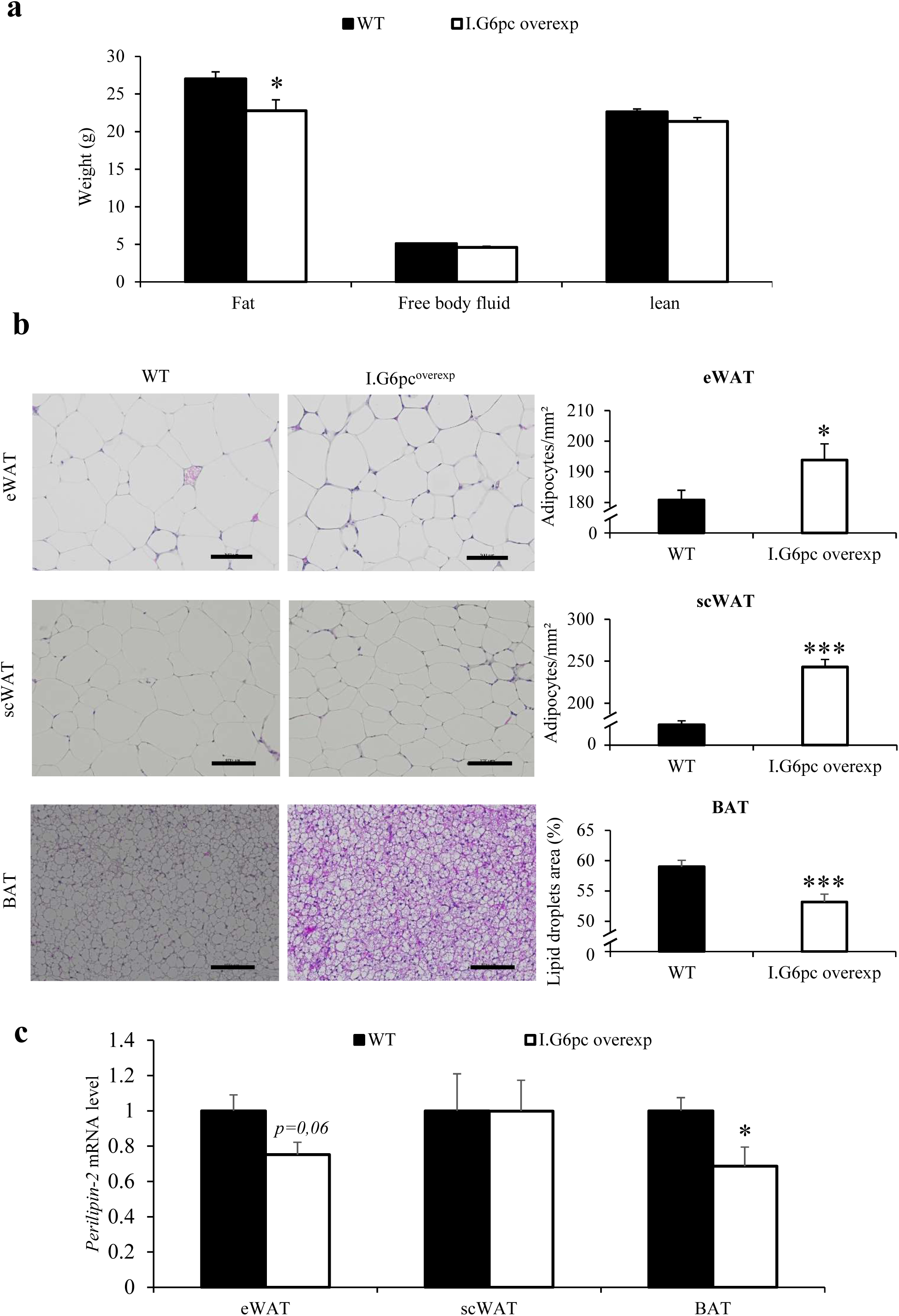
Induction of intestinal gluconeogenesis decreases fat mass expansion induced by high fat-high sucrose diet. **(a)** Body composition of I.G6pc^overexp^ and WT mice (means ± SEM, n= 6-7) **(b)** Representative hematoxylin and eosin staining of eWAT, scWAT and BAT histological sections and quantification (Numbers of adipocytes per mm² for WAT and the lipid droplets area for BAT) (means ± SEM, n= 6; 4-15 pictures per mice). Scale bar represents 100 µm. **(c)** Relative mRNA levels of *Perilipin* in the eWAT, scWAT and BAT (means ± SEM, n= 5). (**a-c)** Student’s t-test was performed as a statistical analysis. *p < 0.05 versus WT mice.

### 3.2. Intestinal gluconeogenesis promotes thermogenesis and prevents whitening processes in brown adipose tissue in mice fed a high fat-high sucrose diet

During the development of obesity, all adipose depots are recruited for lipid storage. In BAT, increased lipid storage leads to its whitening, which refers to the conversion of brown adipocytes in white unilocular cells expressing genes encoding white adipocyte-specific hormones, such as leptin [20,21]. In line with the protection from excessive lipid accumulation described above (**Fig. 1**), *Leptin* expression, a major marker of the whitening of the BAT, was decreased by about 60% in the BAT of I.G6pc^overexp^ mice compared to WT mice (**Fig. 2a)**.

**Figure 2:**
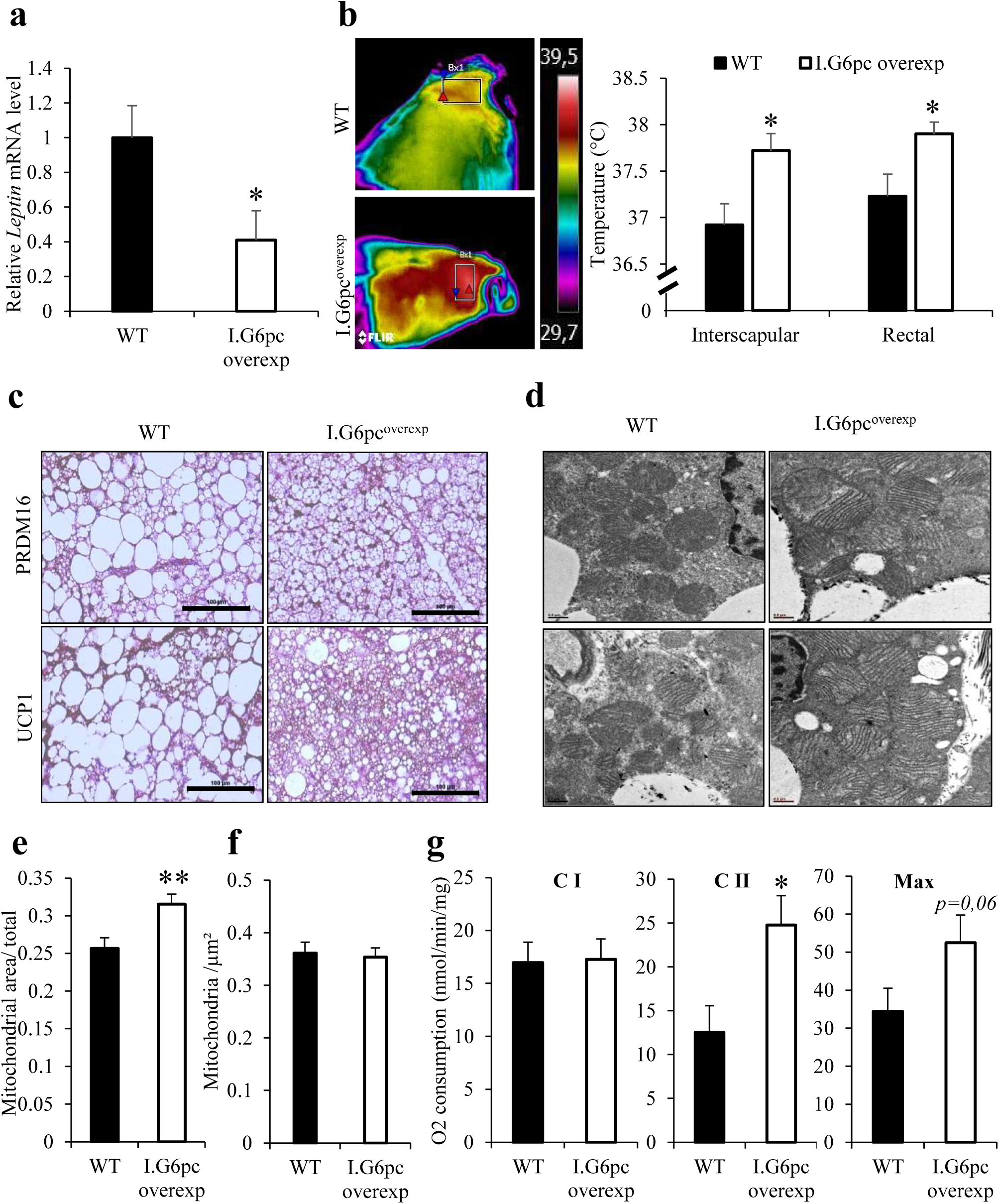
Induction of intestinal gluconeogenesis prevents brown adipose tissue whitening induced by high fat-high sucrose diet. **(a)** Relative mRNA levels of *Leptin* in the BAT (means ± SEM, n= 5-6) **(b)** Skin temperature surrounding the BAT (interscapular region) and rectal temperature (means ± SEM, n= 6-7) **(c)** Representative pictures of UCP1 and PRDM16 staining (in brown) of BAT histological sections. Scale bar represents 100 µm. **(d-e-f)** Representative pictures of electronic microscopy of BAT sections, scale bar represents 0.5 µm (d) and quantification of mitochondria area (e) and number per µm² (f) (n=4; 10 pictures per mice). **(g)** Measure of BAT mitochondria respiratory capacity (means ± SEM, n=12-13). (**a-f)** Student’s t-test was performed as a statistical analysis. *p < 0.05; ** p<0.01 versus WT mice.

While thermogenesis is an essential component of energy expenditure, particularly in the control of energy homeostasis, the loss of BAT thermogenic capacities promotes body weight gain during obesity development [4,22,23]. Interestingly, I.G6pc^overexp^ mice showed a higher inter-scapular and rectal temperature compared to WT mice, suggesting an active thermogenesis taking place in the BAT (**Fig. 2b**). Moreover, we demonstrated by immunochemistry determinations that the amount of two major thermogenic proteins, UCP1 and PRDM16 was higher in the BAT of I.G6pc^overexp^ than in WT mice (**Fig. 2c**). In line with an increase in thermogenesis in the BAT of I.G6pc^overexp^ mice, electron microscopy analysis revealed that mitochondria were larger in these mice compared to WT mice, while the total number of mitochondria was not different (**Fig. 2d-e-f**). This modification in mitochondrial size could impact mitochondrial activity [24,25]. Accordingly, the mitochondrial respiratory capacity was increased at the level of complex II in the BAT of I.G6pc^overexp^ compared to WT mice, with no change in the activity of complex I (**Fig. 2g**). In addition, the experiment using an uncoupling agent (FCCP) demonstrated that there was a trend to a global increase in the maximal mitochondrial respiratory capacity in the BAT of I.G6pc^overexp^ mice (**Fig. 2g**).

Therefore, the induction of IGN prevented the BAT whitening usually occurring under a high calorie diet, thus preserving the BAT thermogenic capacities helping in turn protect from body weight gain.

### 3.3. Intestinal gluconeogenesis promotes the browning of epididymal white adipose tissue in mice fed a high fat-high sucrose diet

White adipose tissue can acquire thermogenic capacities under certain conditions, such as cold exposure, according to a process named “browning”. The browning of WAT is often described as taking place primarily in scWAT during cold exposure [26]. However, analysis of scWAT of I.G6pc^overexp^ mice did not show significant change in the mRNA expression of *Ucp1* and *Prdm16* (**Fig. S1**).

Differently, when analysis was carried out on the eWAT, we observed an increase in the expression levels *Ucp1, Prdm16* in I.G6pc^overexp^ mice (**Fig. 3a**). Interestingly, while the sympathetic nervous system (SNS) is the main regulator of the browning process, the expression level of β*3-adrenergic receptor* (*Adr*β*3*) was increased in I.G6pc^overexp^ mice (**Fig. 3a**). Moreover, the expression of major genes involved in lipid oxidation, as *CoxIV (Cytochrome c oxidase subunit 4), Ppara (Peroxisome proliferator-activated receptor alpha), and the* critical regulator of thermogenesis *Ppargc1a* (*Peroxisome Proliferator-Activated Receptor Gamma Coactivator 1-Alpha*), and in lipolysis, as *Hsl (Hormone sensitive lipase)* were increased in the eWAT of I.G6pc^overexp^, highlighting stimulation of adipocyte metabolism (**Fig. 3b**). However, there was no difference in the size or in the number of mitochondria in the eWAT from both experimental groups (**Fig. 3d-e-f**).

**Figure 3:**
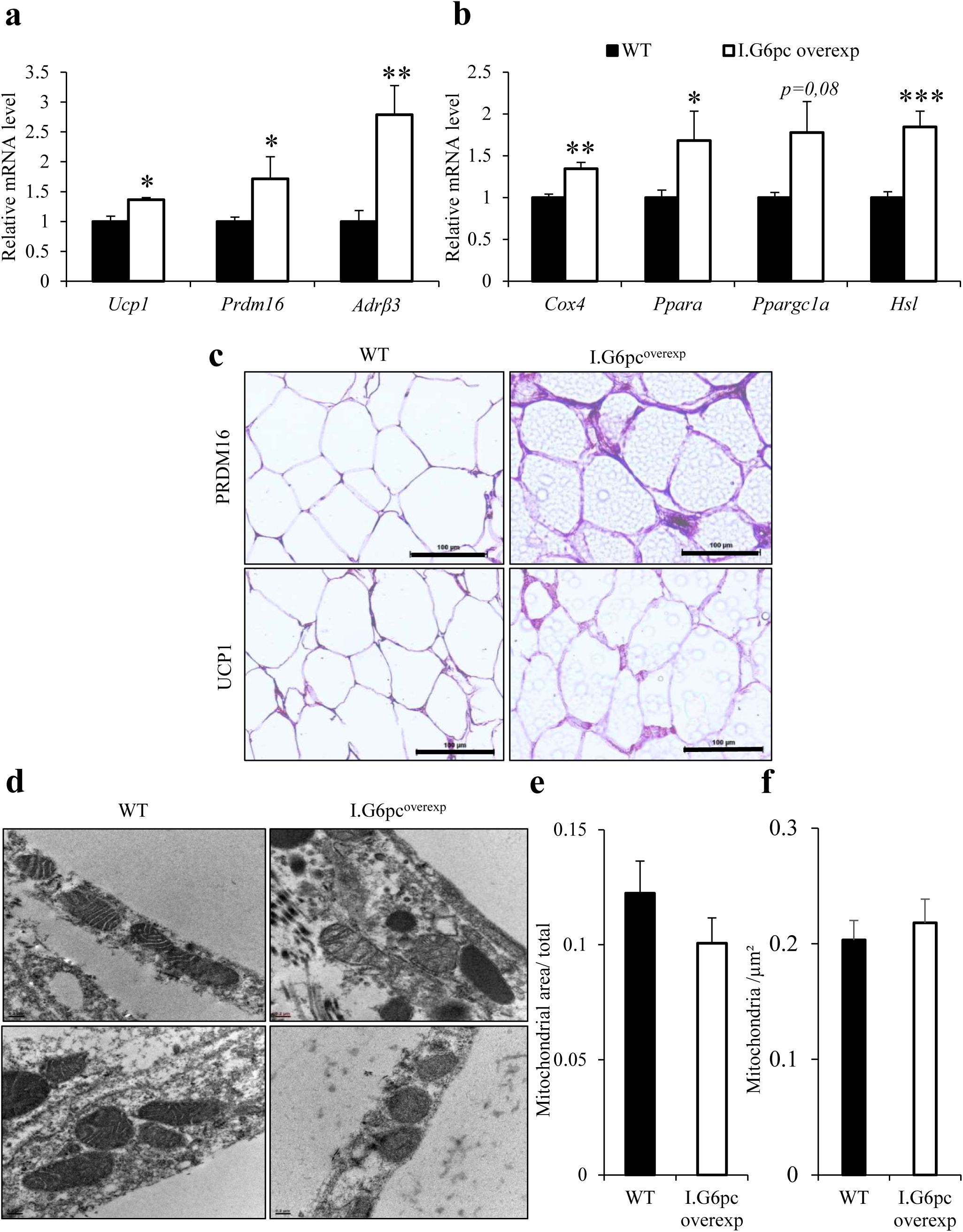
Induction of intestinal gluconeogenesis promotes the browning of epididymal white adipose tissue in mice fed high fat-high sucrose diet. **(a)** Relative mRNA levels of genes implicated in thermogenesis capacity (means ± SEM, n=4-6). **(b)** Relative mRNA levels of genes involved in WAT lipolysis (means ± SEM, n=4-6). **(c)** Representative pictures of UCP1 and PRDM16 staining (in brown) of eWAT histological sections. Scale bar represents 100 µm. **(d-e-f)** Representative pictures of electron microscopy of eWAT sections, Scale bar represents 0.5 µm (d) and quantification of mitochondria area (e) and number per µm² (f) (n=4; 10 pictures per mice). Student’s t-test was performed as a statistical analysis. *p < 0.05; ** p<0.01; ***p<0.001 versus WT mice.

### 3.4. Intestinal gluconeogenesis improves cold resistance by involving the sympathetic nerves

The SNS is the main regulator of BAT thermogenesis [27,28]. Accordingly, increased thermogenesis in I.G6pc^overexp^ mice was associated with a higher protein expression of tyrosine hydroxylase (TH), which is an enzyme essential for the synthesis of catecholamines. (**Fig. 4a**).

**Figure 4:**
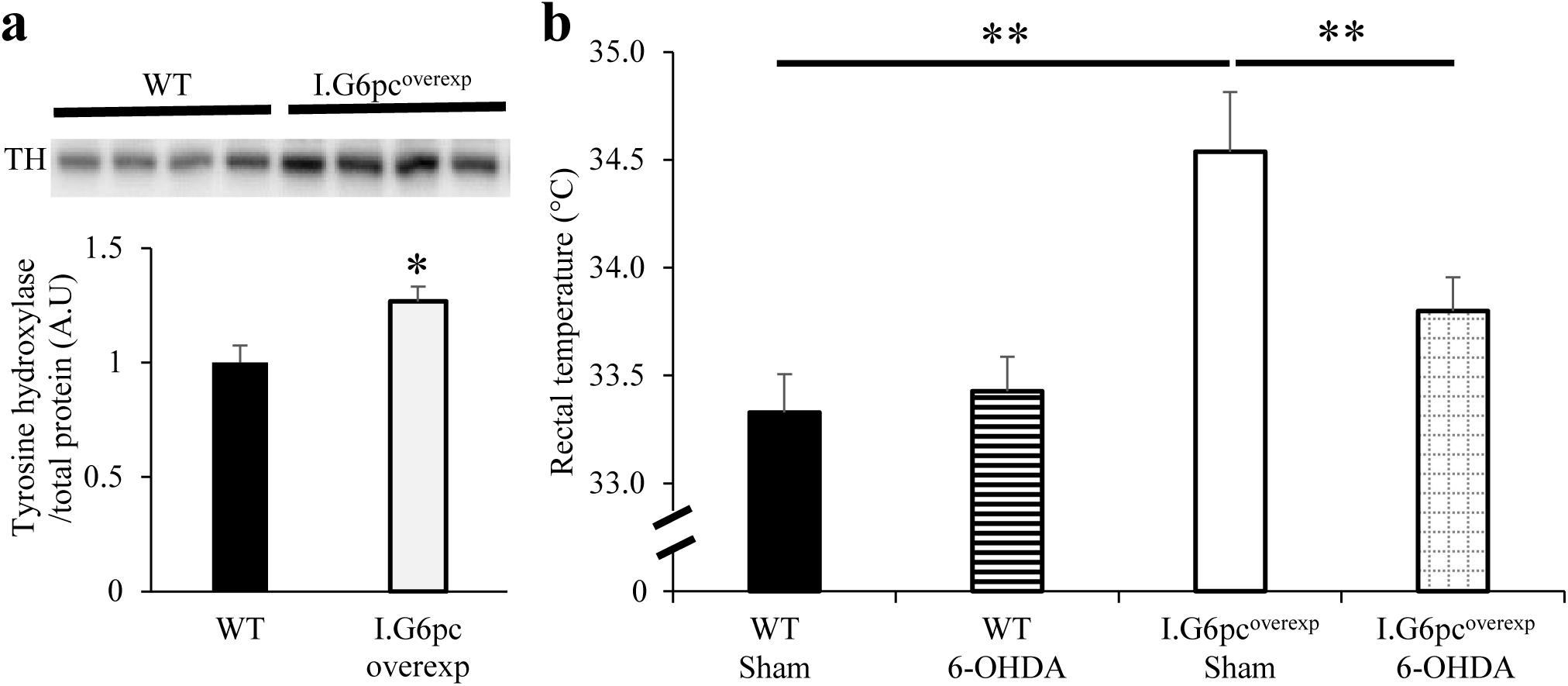
Induction of intestinal gluconeogenesis potentiates the resistance to cold and increases energy expenditure in mice fed high fat-high sucrose diet. **(a)** Relative quantification of tyrosine hydroxylase protein levels in the eWAT (means ± SEM, n=4-6) and representative western blot picture. **(b)** Rectal temperature of mice after sympathetic denervation restricted to the BAT (6-OHDA-treated vs Sham) following 4-hours cold exposure (6°C) (means ± SEM, n=7-8). **(a-b)** Student’s t-test (A) or ANOVA-2 followed by Bonferroni post-hoc test (B) was performed as a statistical analysis. *p < 0.05; ** p<0.01 versus WT mice.

To further investigate the effects of IGN in the regulation of adipose tissue thermogenesis, we studied the effects of its activation on cold resistance in a standard dietary context. More specifically, we investigated whether the increase in thermogenesis in I.G6pc^overexp^ mice (Figures 2-3 above) could confer on them a better resistance to cold even on a standard chow diet. In addition, we tested the involvement of the SNS in the transmission of the beneficial effects of IGN relating to the resistance to cold. For this purpose, we performed injections restricted to the BAT of a specific SNS neurotoxin, 6-OHDA. Ten-days after the surgery, WT and I.G6pc^overexp^ mice, subdivided into two groups - control (sham/vehicle treated) and denervated (6-OHDA treated) - were exposed to cold and their rectal temperature measured (**Fig. 4b**). While we observed the improvement of cold resistance in I.G6pc^overexp^-sham mice, these effects did not take place in the absence of sympathetic nerve fibres in the BAT in I.G6pc^overexp^ 6-OHDA treated mice (**Fig. 4b**). Thus, while IGN increases thermogenesis both in the BAT and eWAT, these data suggest a prominent role for BAT in maintaining body temperature during short cold exposure (4-hours; 6°C) (**Fig. 4b**). In addition, these results were not accompanied by a change in food intake, which excludes any interference of diet-induced thermogenesis with the effects (**Fig. S2**).

Therefore, while the activation of IGN increased thermogenesis, using a chemical SNS denervation approach, we clarified that the modulation of SNS is necessary to observe this modulation deriving from IGN activation.

### 3.5. Intestinal gluconeogenesis modulates adipokine secretion in the context of high fat-high sucrose diet

During the development of obesity, the WAT is a site of numerous deleterious changes, including the increase in the production and release of proinflammatory cytokines. The later can lead to the proliferation of the extra-cellular matrix, contributing to fibrosis development in adipose tissue [29,30]. Interestingly we observed that the eWAT of I.G6pc^overexp^ mice showed a decrease in the gene expression of the proinflammatory marker TNFα (Tumor necrosis factor alpha), without significant change in the expression of other proinflammatory adipokines (**Fig. 5a**). In addition, the observation of Masson trichrome-stained eWAT sections strongly suggested a prevention of fibrosis in I.G6pc^overexp^ mice (**Fig. 5b**)

**Figure 5:**
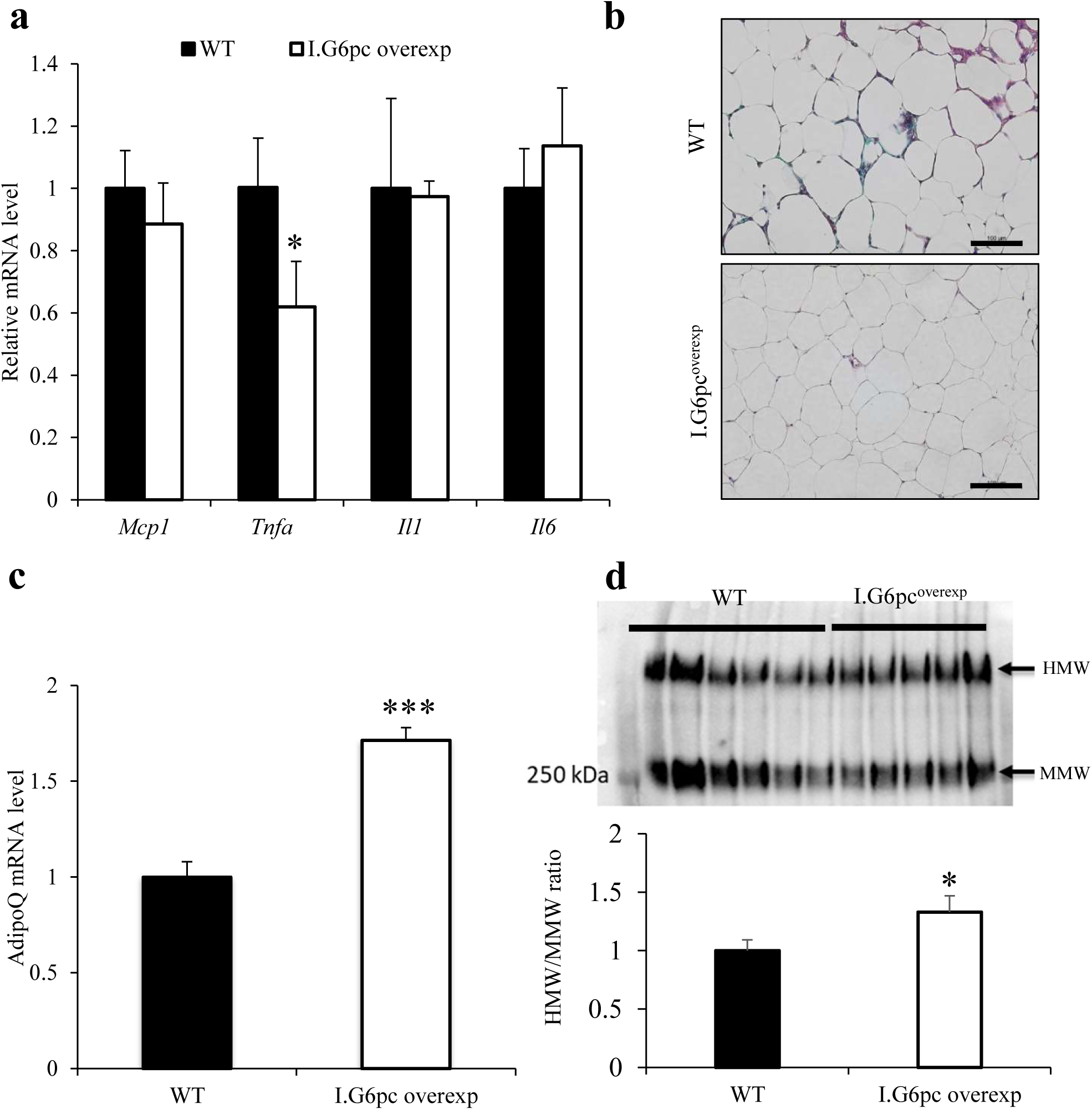
Induction of intestinal gluconeogenesis protects against deleterious changes in adipokine secretion profile induced by high fat-high sucrose diet. **(a)** Relative mRNA levels of genes involved in pro-inflammatory processes (means ± SEM, n=5-6). **(b)** Masson trichrome staining of liver histological sections. Scale bar represents 100 µm. **(c)** mRNA levels of *AdipoQ* expression (means ± SEM, n=4-6). **(d)** Representative image of non-denaturing SDS-PAGE in the eWAT. The graph shows the quantification of High Molecular Weight (HMW)/Medium Molecular Weight (MMW) forms (means ± SEM, n= 10-12). **(a-d)** Student’s t-test was performed as a statistical analysis. *p < 0.05; ***p<0.001 versus WT mice.

In parallel, we observed an increase in the mRNA expression of adiponectin *(AdipoQ)* (**Fig. 5c**), an adipokine known to improve insulin sensitivity, lipid metabolism and inflammation [31]. Moreover, we measured an increase in the high molecular weight form (HMW) of adiponectin, which corresponds to the biologically active form exerting the majority of its beneficial metabolic effects [32] (**Fig. 5d**).

### 3.6. Specific suppression of intestinal gluconeogenesis promotes lipid storage and alteration in adipokine secretion in mice fed a standard diet

To get further insight in the mechanisms by which IGN modulates the metabolism of BAT and WAT, we used mice with specific deletion of IGN (I.G6pc^-/-^ mice). At the opposite of what occurred in I.G6pc^overexp^ mice, the absence of this function promoted lipid storage in the WAT, illustrated by an increase in body weight (**Fig. 6a**) and in the weight of the eWAT and scWAT in I.G6pc^-/-^ mice compared with WT mice (**Fig. 6b-c**). Hematoxylin-stained sections of adipose tissues revealed a hyperplasia of adipocytes in the eWAT and scWAT and an increased area occupied by lipid droplets in the BAT of I.G6pc^-/-^ mice, compared to WT mice (**Fig. 6d**).

**Figure 6:**
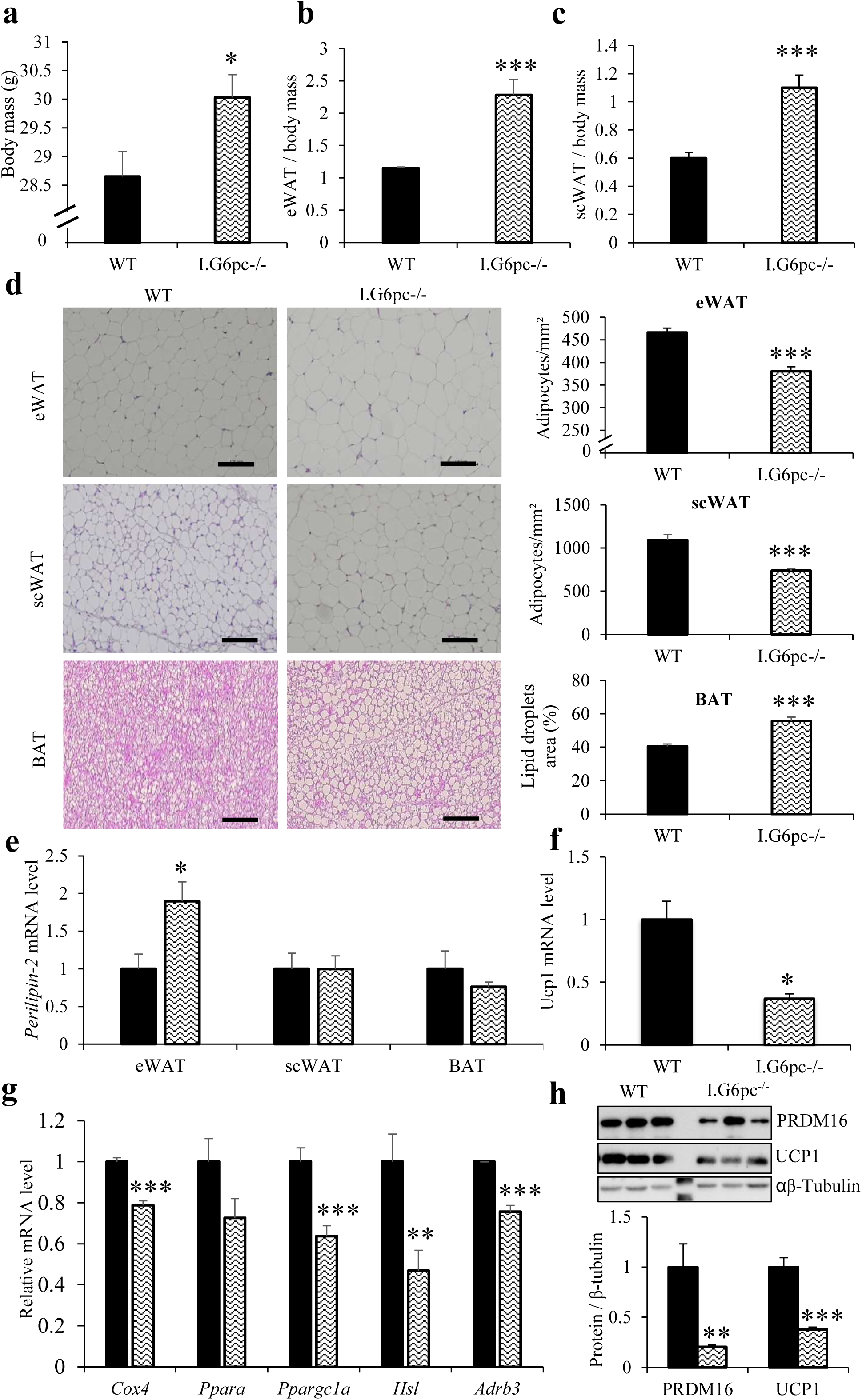
Absence of intestinal gluconeogenesis promotes adipose tissue alterations (in standard diet) **(a)** Body mass of I.G6pc^-/-^ and WT mice (means ± SEM, n=7-8). **(b-c)** eWAT and scWAT mass relative to body mass (means ± SEM, n=7-8). **(d)** Representative hematoxylin and eosin staining of eWAT, scWAT and BAT histological sections and quantification (Numbers of adipocytes per mm² for WAT and the lipid droplets area for BAT) (means ± SEM, n= 6; 4-15 pictures per mice). Scale bar represents 100 µm. **(e)** Relative *Perilipin* mRNA expression levels in the eWAT, scWAT and BAT (means ± SEM, n=4-6). **(f)** Relative mRNA levels of *Ucp1* in the BAT (means ± SEM, n=3-4). **(g)** Relative mRNA levels of genes involved in eWAT lipolysis and lipid oxidation (means ± SEM, n=4-6). **(h)** Relative quantification of protein levels of PRDM16 and UCP1 in the eWAT (means ± SEM, n=5) and representative western blot picture. (**a-h)** Student’s t-test was performed as a statistical analysis. *p < 0.05; ***p<0.001 versus WT mice.

In agreement with increased adiposity in I.G6pc^-/-^ mice, *Perilipin* expression was increased in the eWAT, albeit not in the scWAT and BAT (**Fig. 6e**). Relating to thermogenic markers, the BAT of I.G6pc^-/-^ mice exhibited a decrease in the expression of *Ucp1* (**Fig. 6f**).

It is noteworthy that, at the opposite of what took place in I.G6pc^overexp^ mice, IGN deficiency led to a decrease in the expression of genes involved in thermogenic capacities, such as *Ppargc1a, CoxIV* and *Hsl* in the eWAT (**Fig. 6g**). In agreement, the expression level of the thermogenesis markers UCP1 and PRDM16 were also decreased in the eWAT of I.G6pc^-/-^ mice (**Fig. 6h**). In addition, we showed a decrease in *Adrb3* expression levels (**Fig. 6g**).

Interestingly, the absence of IGN was sufficient to increase the expression level of the pro-inflammatory markers *Il-1* and *Il-6*, and the necrosis factor *Tnfa* in the eWAT (**Fig. 7a-b**). Furthermore, in mirror again of the situation in I.G6pc^overexp^ mice, IGN deficiency lead to a decrease in the expression level of *AdipoQ* and of the HMW conformation of Adiponectin. (**Fig. 7c-d**).

**Figure 7:**
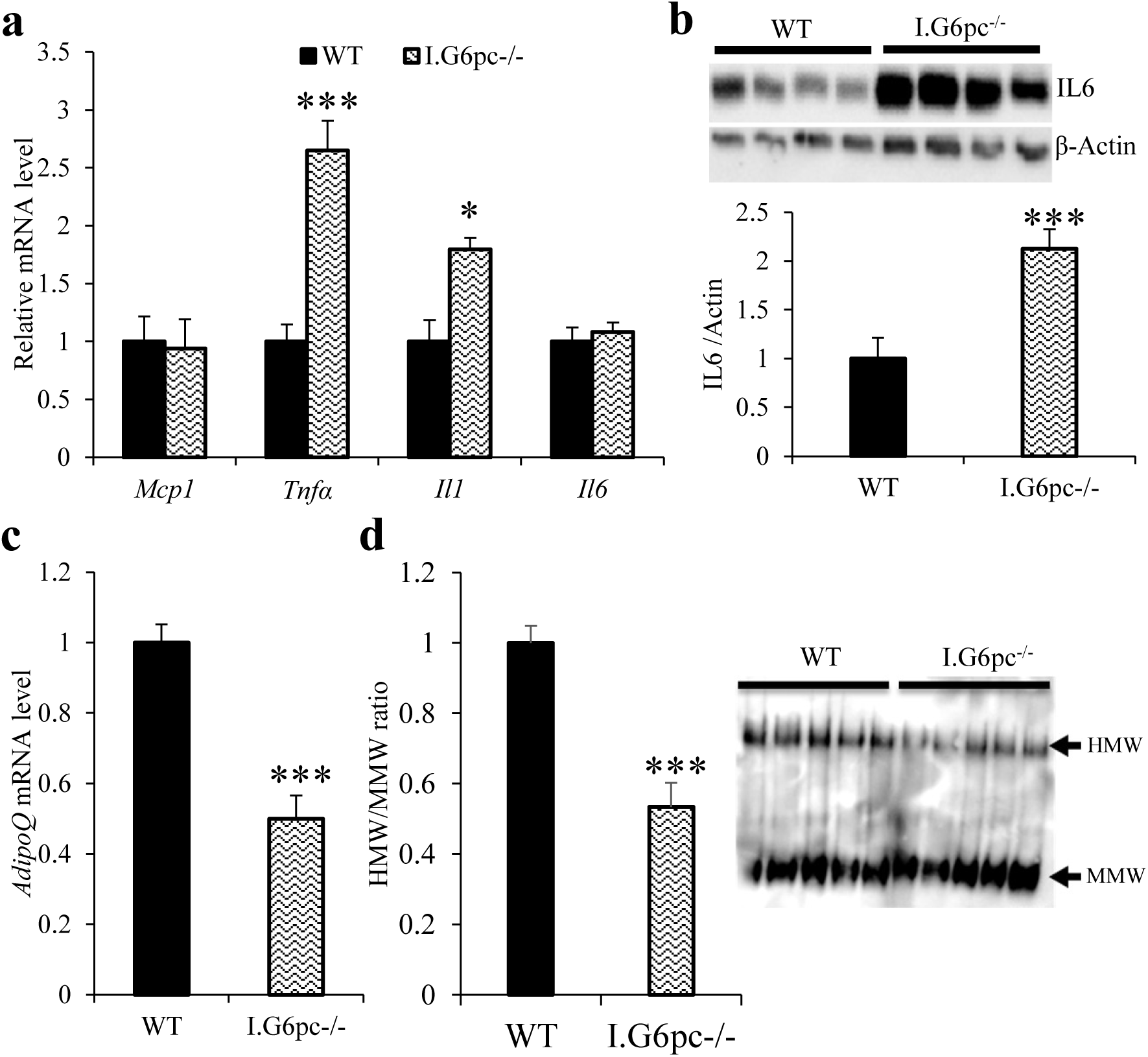
Absence of intestinal gluconeogenesis promotes deleterious changes in adipokine secretion profile (in standard diet) **(a)** Relative mRNA levels of genes involved in pro-inflammatory processes (means ± SEM, n=5-6). **(b)** Relative quantification of protein levels of IL6 in the eWAT (means ± SEM, n=5) and representative western blot picture. **(c)** Relative mRNA levels of *AdipoQ* expression (means ± SEM, n=7). **(d)** Representative image of non-denaturing SDS-PAGE in the eWAT. The graph shows the quantification of MHW/MMW forms (means ± SEM, n= 5). **(a-d)** Student’s t-test was performed as a statistical analysis. *p < 0.05; ***p<0.001 versus WT mice.

Altogether, these data highlight that the absence of IGN is sufficient to alter adipose tissue metabolism, even in young mice (13-weeks old) and in a standard feeding context.

## 4. DISCUSSION

The metabolic benefits conferred by the induction of IGN were previously observed upon various nutritional manipulations. Indeed, IGN is a causal link in the anti-obesity and anti-diabetic effects of dietary proteins and soluble fibres and of gastric bypass surgery [7,8,10,11,13,14,33]. In order to study the beneficial effects of IGN *per se*, independently of the nutritional context, we used mouse models of overexpression or suppression of IGN, by targeting G6PC, the catalytic subunit of G6Pase.

In agreement with a marked effect of prevention of obesity, the induction of IGN leads to a reduction of lipid storage in eWAT, scWAT and BAT. The expansion of adipose tissue during the development of obesity is associated with deregulation of fatty acid fluxes and can lead to hypoxia and adipocyte death, which promotes pro-inflammatory and insulin-resistant states (for review see [34]). The prevention of this expansion by IGN is therefore a key benefit in the context of metabolic diseases.

The BAT is the main player in non-shivering thermogenesis. In obese subjects, BAT undergoes profound deleterious changes. Especially, the activity of BAT decreases while adiposity increases [35,36]. Thus, BAT thermogenesis represents a therapeutic target in the treatment of obesity and the prevention of its associated complications [37,38]. In this context, we show here that I.G6pc^overexp^ mice exhibit an increase in heat production by the BAT associated with higher body temperature. Moreover, our data strongly suggest that the induction of IGN protects the BAT against the whitening process known to occur in deleterious nutritional contexts (such as high calorie diet) and to induce the loss of thermogenic capacities [4,22,23].

The morphology of mitochondria is a key parameter for an optimal metabolic activity of BAT, especially for thermogenesis [25,37]. Accordingly, mitochondria in the BAT are more numerous and larger than their counterparts in the WAT [24]. Therefore, the increase in mitochondria size in the BAT of I.G6pc^overexp^ mice, associated with increased expression of UCP1 and PRDM16, could contribute to the induction of heat production. Moreover, the induction of UCP1-dependent thermogenesis by cold exposure requires an increase in the activity of complex II of the respiratory chain, essential for the oxidation of FADH2 (Dihydroflavine-adenine dinucleotide) generated from lipid oxidation, while an inhibition of complex II suppresses cold-induced thermogenesis [39]. Accordingly, the increase in complex II activity measured in the BAT of I.G6pc^overexp^ mice is in agreement both with the increased heat production by the interscapular region and with the associated mobilization of lipids for that.

Both BAT thermogenesis and the process of browning in WAT, which corresponds to the appearance of beige adipocytes capable of producing heat, are controlled by the SNS [37,38,40,41]. In line with this, IGN induction led to an increase in the expression levels of thermogenic markers and β3-adrenergic receptor in the eWAT. In addition, the protein expression level of tyrosine hydroxylase, an enzyme essential for the synthesis of catecholamines, was increased in the BAT of I.G6pc^overexp^ mice fed HF-HS diet. Interestingly, while this increased thermogenesis in both BAT and WAT likely contributes to both the better resistance to cold in I.G6pc^overexp^ mice, we clarify here the mechanisms evoked by the activation of the IGN in adipose tissues from a SNS chemical denervation approach. Indeed, this effect doesn’t occur after the inactivation by 6-OHDA of BAT-sympathetic innervation. Thus, although the increased WAT browning can be beneficial in maintaining energy balance on HF-HS diet, the effect of IGN on BAT thermogenesis appears to be predominant compared with that in WAT regarding thermogenesis and cold resistance. Altogether, these data strongly suggest that IGN promotes the BAT thermogenesis and the browning of the WAT by involving a gut-brain-adipose tissue neural loop activating the sympathetic tone.

It is noteworthy that mice lacking IGN show a picture in mirror compared to I.G6pc^overexp^ mice, including an increase in the size of adipocytes and lipid droplets associated with a decrease in the expression levels of thermogenic markers in both the eWAT and BAT. These alterations could be linked to the development of dyslipidaemia observed in the absence of IGN [16]. Indeed, the decrease in adipose tissue thermogenesis could be responsible, at least in part, for the increase in plasma triglycerides observed in I.G6pc^-/-^ mice [16]. This excess may contribute to ectopic lipid accumulation in others tissues, particularly in the liver, thus contributing to worsen hepatic insulin resistance in I.G6pc^-/-^ mice [16]. Conversely, the activation of both the eWAT and BAT metabolism in I.G6pc^overexp^ mice could contribute to the prevention of hepatic lipid storage and the moderation of body weight gain on HF-HS diet [16].

Interestingly, the adipose tissue is also an important endocrine tissue implicated in the regulation of carbohydrate and energy homeostasis *via* the secretion of the so-called adipokines [42]. During the development of obesity, the secretion profile of these hormones is altered with a decrease in the beneficial adipokines, such as adiponectin, in favour of pro-inflammatory adipokines [42]. A decrease in circulating adiponectin levels in obese patients was proposed to have a crucial role in the pathogenesis of metabolic complications associated with obesity, especially hepatic steatosis [43,44]. It is noteworthy that I.G6pc^overexp^ mice show an increase in gene expression of adiponectin and of its HMW conformation that is the main biologically active form mediating its metabolic benefits, such as the improvement of hepatic insulin sensitivity or of lipid metabolism [43,44]. Thus, these changes in adiponectin metabolism may be involved in the protection against the development of hepatic steatosis and the observed delay in the onset of diabetes on high calorie diet in I.G6pc^overexp^ mice [16]. In contrast, I.G6pc^-/-^ mice exhibit a decrease in the HMW adiponectin form. These alterations could be partly involved in the development of the pre-diabetic state and increase in hepatic lipid content previously observed in I.G6pc^-/-^ mice, notably characterized by hepatic insulin resistance [16,45].

It is noteworthy that several studies suggested functional differences in white adipose tissue depots depending on their anatomical location. While the mass of visceral adipose tissue is associated with peripheral insulin resistance, the preferential accumulation of lipids in the scWAT could be protective against metabolic complications [46,47]. Accordingly, the surgical removal of eWAT in rodents prevents the development of obesity, insulin resistance, dyslipidaemia and hepatic steatosis while increasing the storage in the scWAT, a process possibly linked to adipokines [47]. Thus, it seems particularly interesting on a physiological viewpoint that IGN increases browning in a visceral depot as the eWAT, since, in addition, this fat pad is generally suggested to be less susceptible to browning compared to the scWAT [48–50].

## 5. CONCLUSION

In conclusion, we here decipher the beneficial effects of IGN in the regulation of energy homeostasis. IGN positively controls fat storage, thermogenic capacities and the adipokine secretion profile through a gut-brain-adipose tissue neural circuit involving the sympathetic nerves. Moreover, this modulation may take place from the only activation of IGN, independently of any additional nutritional manipulation. Therefore, these data highlight that activating IGN constitutes a potential therapeutic target to prevent or correct obesity and its deleterious metabolic consequences, including the pathological expansion of adipose tissue and the thermogenesis decline. This could account at least in part for the weight loss processes occurring after gastric bypass surgery or upon fibre feeding in obese humans. Thus, these data may pave the way to novel approaches of prevention or treatment of metabolic diseases as obesity or type-2 diabetes. The search for molecules capable of activating the IGN and its beneficial metabolic effects is a major challenge for the coming years.

## ABBREVIATIONS

AdipoQ: Adiponectin gene
ADP: Adenosine diphosphate
Adrβ3: β3-adrenergic receptor
BAT: Brown adipose tissue
CoxIV: Cytochrome c oxidase subunit 4
eWAT: Epididymal white adipose tissue
FADH2: Dihydroflavine-adenine dinucleotide
FCCP: p-trifluoromethoxy-carbonyl-cyanide-phenyl hydrazone
G6Pase: Glucose-6-phosphatase
G6PC: Catalytic subunit of Glucose-6-phosphatase
HF-HS: Hgh fat-high sucrose
HMW: high molecular weight form of adiponectin
Hsl: Hormone sensitive lipase
IGN: Intestinal gluconeogenesis
MRI: Magnetic resonance imaging
6-OHDA: 6-hydroxydopamine
Ppara: Peroxisome proliferator-activated receptor α
Ppargc1a: Peroxisome Proliferator-Activated Receptor Gamma Coactivator 1-α
PRDM16: PR domain containing 16
PVDF: Polyvinylidene fluoride
scWAT: Subcutaneous white adipose tissue
SDS: Sodium dodecyl sulfate
SEM: Standard error of the mean
SGLT3: Sodium-glucose co-transporter 3
SNS: Sympathetic nervous system
TH: Tyrosine hydroxylase
TNFα: Tumor necrosis factor α
UCP1: Uncoupling Protein 1
WAT: White adipose tissue
WT: Wild-type

## ACKNOWLEDGMENTS

We thank the members of Animalerie Lyon Est Conventionnelle et SPF (ALECS, Université Lyon 1, SFR Santé Lyon Est) for animal housing and care, and the members of the CIQLE platform (Université Lyon 1, SFR Santé Lyon Est) for histological and electron microscopy experiments (Université Lyon 1).

## AUTHOR CONTRIBUTIONS

J.V-P. designed, conducted experiments, analysed the data and wrote the paper. M.S-R. designed, conducted experiments and analysed the data. M.S. M.M, C.B and M.R. were in charge of mouse breeding and animal experiments. D.B and R.N performed MRI and skin temperature surrounding BAT measurement and analysis. D.R performed the measurement and the analysis of mitochondria respiratory capacity. R.N, D.R, A.G-S, F.R and D.C critically edited the manuscript. G.M. supervised the studies and wrote the paper

## FUNDING STATEMENT

This work was supported by research grants from the Institut National de la Santé et de la Recherche Médicale (INSERM), the Agence Nationale de la Recherche (ANR-17-CE14-0020-01), the Fondation pour la Recherche Médicale (Equipe FRM DEB20160334898) and the company Servier.

## CONFLICT OF INTEREST STATEMENT

None of the authors has a conflict of interest to disclose relating to the work described in this paper.

---

**Supplemental figure 1:**
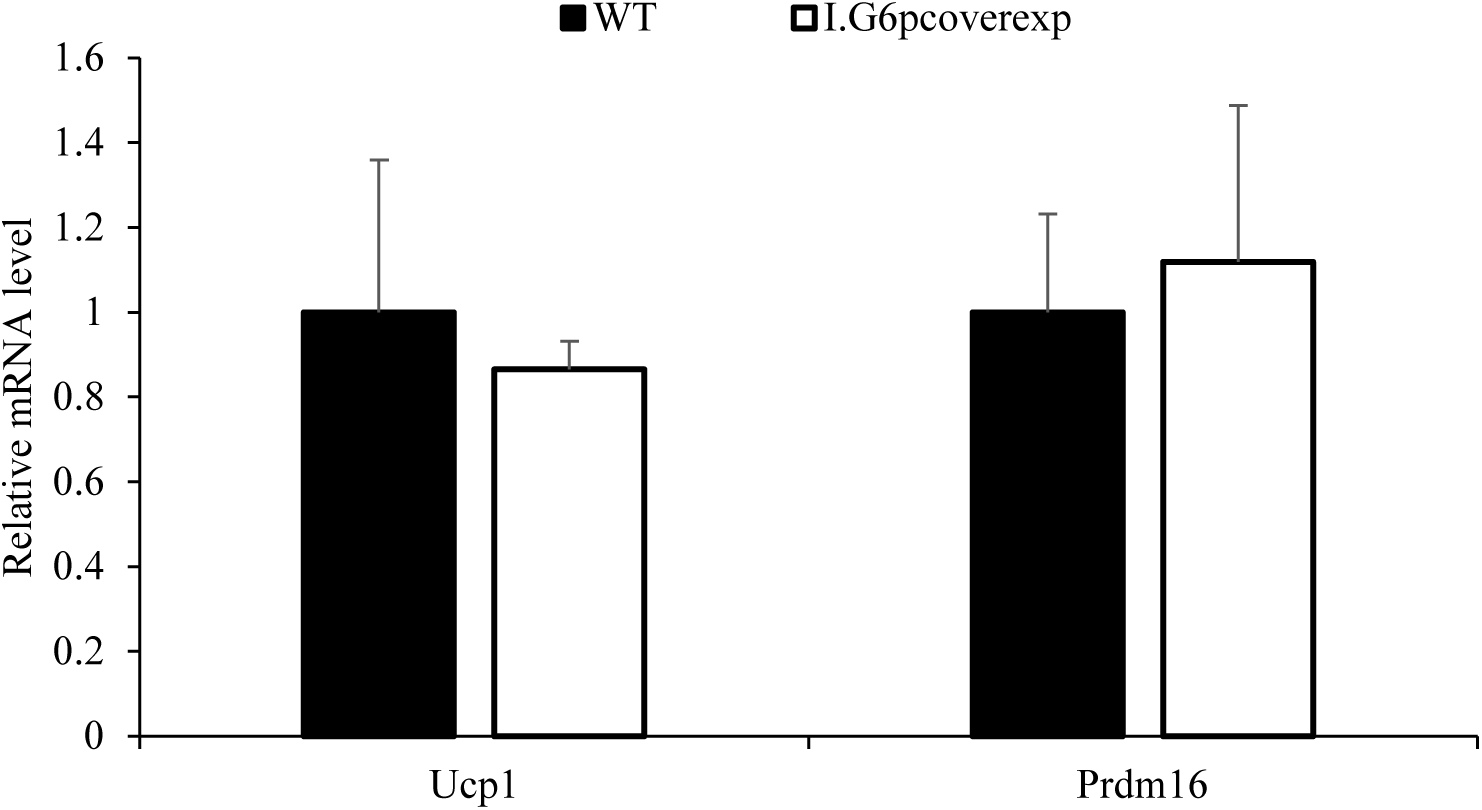
RNAm expression level of thermogenesis markers in subcutaneous white adipose tissue of I.G6pc^overexp^ and WT mice. Student’s t-test was performed as a statistical analysis (means ± SEM, n=4-6).

**Supplemental figure 2:**
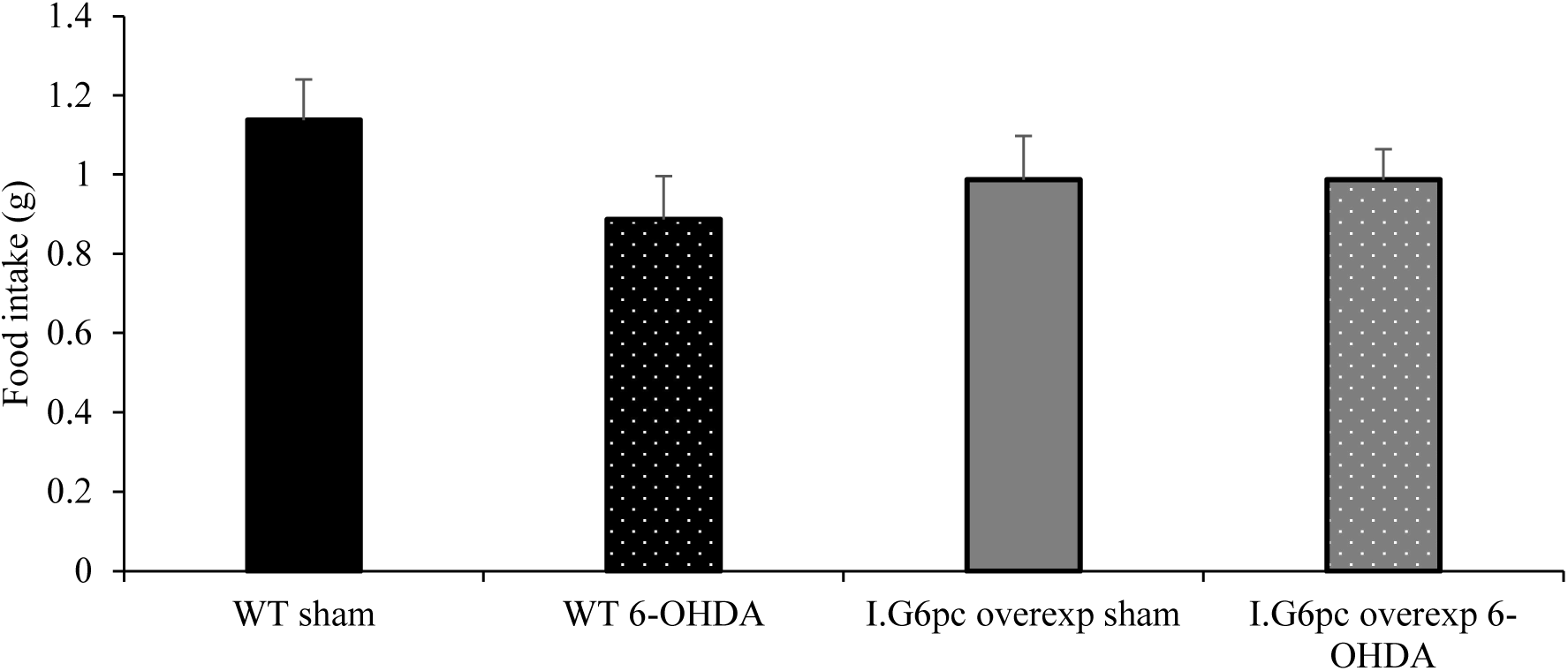
Effects of denervation of the sympathetic system innervating brown adipose tissue on food intake of WT and I.G6pc^overexp^ mice during 4 hours of cold exposure. (means ± SEM, n=6-7). Student’s t-test was performed as a statistical analysis.

## Notes

### Competing Interest Statement

The authors have declared no competing interest.

